# Germline Cas9 Expression Yields Highly Efficient Genome Engineering in a Major Worldwide Disease Vector, *Aedes aegypti*

**DOI:** 10.1101/156778

**Authors:** Ming Li, Michelle Bui, Ting Yang, Bradley J. White, Omar S. Akbari

**Author notes:** Corresponding Author: Omar S. Akbari, 4484 Boyce Hall, Riverside CA, 92521.

## Abstract

The development of CRISPR/Cas9 technologies has dramatically increased the accessibility and efficiency of genome editing in many organisms. In general, *in vivo* germline expression of Cas9 results in substantially higher activity than embryonic injection. However, no transgenic lines expressing Cas9 have been developed for the major mosquito disease vector *Aedes aegypti*. Here, we describe the generation of multiple stable, transgenic *Ae. aegypti* strains expressing Cas9 in the germline, resulting in dramatic improvements in both the consistency and efficiency of genome modifications using CRISPR. Using these strains, we disrupted numerous genes important for normal morphological development, and even generated triple mutants from a single injection. We have also managed to increase the rates of homology directed repair by more than an order of magnitude. Given the exceptional mutagenic efficiency and specificity of the Cas9 strains we built, they can be used for high-throughput reverse genetic screens to help functionally annotate the *Ae. aegypti* genome. Additionally, these strains represent a first step towards the development of novel population control technologies targeting *Ae. aegypti* that rely on Cas9-based gene drives.

**Significance Statement:** *Aedes aegypti* is the principal vector of multiple arboviruses that significantly affect human health including dengue, chikungunya, and zika. Development of tools for efficient genome engineering in this mosquito will not only lay the foundation for the application of novel genetic control strategies that do not rely on insecticides, but will also accelerate basic research on key biological processes involved in disease transmission. Here, we report the development of a transgenic CRISPR approach for rapid gene disruption in this organism. Given their high editing efficiencies, the Cas9 strains we developed can be used to quickly generate novel genome modifications allowing for high-throughput gene targeting, and can possibly facilitate the development of gene drives, thereby accelerating comprehensive functional annotation and development of innovative population control strategies for *Ae. aegypti*.

## Introduction

The yellow fever mosquito*, Aedes aegypti,* is the principal vector of many arboviruses such as dengue, chikungunya, yellow fever, and zika. These pathogens are globally widespread and pose significant epidemiological burdens on infected populations, resulting in hundreds of millions infections and over 50,000 deaths per year (1–4). Due to the hazards they impose, many methods for controlling *Ae. aegypti* populations have been implemented with the most common being chemical insecticides. However, chemical control has proven incapable of stopping the spread of *Ae. aegypti,* primarily due to its ability to rapidly adapt to new climates, tendency to oviposit in minimal water sources, desiccation-tolerant eggs, and quick development of insecticide resistance (5, 6). Therefore, significant efforts are currently underway to discern the underlying molecular and genetic mechanisms important for arboviral vector competence with the overall aim of developing novel insecticide-free ways to disrupt viral disease cycles (7).Importantly, uncovering these mechanisms hinges on the ability to stably insert and disrupt specific genes-of-interest through tailored genome engineering in a target specific manner. Fortunately, several tools have been successfully employed in mosquitoes for targeted genome engineering that rely on either zinc finger nucleases (ZFN’s)(8–10), transcription activator-like effector nucleases (TALEN’s)(11, 12), or even homing endonuclease genes (HEG’s)(13).However, because they rely on context-sensitive modular protein-DNA-binding interactions, each of these designer nucleases are time-consuming and complicated to engineer and validate, making them onerous for routine use in most labs.

To overcome the significant limitations posed by previous genome editing tools, the clustered regularly interspaced short palindromic repeats/CRISPR-associated sequence 9 (CRISPR/Cas9) system, originally discovered in bacteria and archaea (14–20), has been adapted as a programmable (20, 21) precision genome editing tool in a diversity of organisms (22–29), including mosquitoes (30–34). Briefly, the CRISPR/Cas9 system is guided by a chimeric programmable synthetic short guide RNA (sgRNA) (20) that binds Cas9 directing it to a user specified genomic DNA target sequence, via Watson-Crick base pairing between the sgRNA and the target DNA sequence, thereby generating site-specific double-strand (ds) dsDNA breaks.This system can be easily reprogrammed to modify virtually any desired genomic sequence by recoding the specificity-determining sequence of the sgRNA. Recoding constraints are minimal and simply require that the target sequence is unique compared to the rest of the genome, and is located just upstream of a protospacer adjacent motif (PAM sequence), typically consisting of the 3 nucleotide (nt) motif NGG (20, 21). Both the minimal constraints and ease of use make CRISPR/Cas9 a powerful tool for genome engineering applications.

Importantly, CRISPR-mediated genome engineering of organisms has been achieved in a variety of different ways. For example, direct injection of *in vitro* purified sgRNAs combined with either purified Cas9 RNA, recombinant Cas9 protein, or even with Cas9 expression plasmids, have all been successful for a variety of organisms (29, 32, 35–40), including mosquitoes (30–34). However, the rate of mutagenesis and lethality have varied widely both within and among these different studies, and such discrepancies are likely due to either the methods used to introduce the editing components, the variability in sgRNA functionality against target sites, or even unavoidable variability from manual injection of the components (41). To overcome these significant limitations, previous studies in other organisms have shown that stably expressing a transgenic provision of Cas9 in the germline can decrease toxicity to injected embryos, increase the rates of mutagenesis generated by both non-homologous end joining (NHEJ) and homology directed repair (HDR), and can also increase the rates of germline transmission of the disrupted allele to offspring (27, 41–46).

Germline expression of Cas9 is also essential for developing innovative technologies that rely on “gene drive,” which is a novel strategy proposed for the control of vector-borne diseases by rapidly spreading alleles in a population through super-Mendelian inheritance (47, 48). In mosquitoes, gene drives could potentially be used to rapidly disseminate a genetic payload that reduces pathogen transmission throughout a population, thereby suppressing vector competence and human disease transmission. Other possible applications include the suppression of the population by spreading alleles that impair fertility or viability (Burt 2003; Deredec, Burt, and Godfray 2008). A CRISPR homing based gene drive element consists of only a few components such as an sgRNA and a germline expressed Cas9 endonuclease that is positioned opposite its target site in the genome. The drive encodes the editing machinery (i.e. Cas9 and sgRNA) allowing it to cut the opposite allele and copy itself into this disrupted allele via HDR, thereby converting a heterozygote into a homozygote, enabling rapid invasion of the drive into a population. In fact, Cas9 has been used to develop highly promising homing based gene drives in a number of organisms including yeast (49), *Drosophila* (50), and *Anopheles* mosquitoes (50, 51), however such a system has yet to be developed in *Ae. aegypti*.

We therefore aimed to develop a CRISPR/Cas9 transgenic system that would enable more robust and widespread *Ae. aegypti* genome engineering applications including disrupting genes important for vector competence, while also laying the foundation for the future development of gene drives. To do so, we utilized several previously described transcriptional regulatory elements (52, 53), many of which are active in the germline (54), to drive expression of *S. pyogenes* Cas9 for the first time in *Ae. aegypti*. In total we develop six independent Cas9 expressing strains, and characterize and demonstrate their effectiveness for gene disruption, and gene insertion via HDR. We demonstrate the efficiency of using these strains by disrupting six novel genes in *Ae. aegypti*, giving rise to severe phenotypes such as having an extra eye (triple eyes), an extra maxillary palp (triple maxillary palps), a non-functional curved proboscis, malformed wings, eye pigmentation deficiencies, and pronounced whole animal cuticle coloration defects. Furthermore, we also demonstrate the ease of generating double and triple mutant strains simultaneously from a single injection, a technique that will facilitate the ease of gene function study in this non-model organism. Overall, gene disruption efficiencies, survival rates, germline transmission frequencies, and HDR rates were all significantly improved using the Cas9 strains we develop here. These strains should be highly valuable for facilitating the development of innovative control methods in this organism in the future.

## Results

### Construction of a simple transgenic CRISPR/Cas9 system for *Ae. aegypti* mutagenesis

In order to express Cas9 in the germline of *Ae. aegypti*, we established transgenic mosquitoes harboring genomic sources of Cas9. To promote robust expression of Cas9, we utilized promoters from six genes including AAEL010097 (*Exuperentia*), AAEL007097 (4-nitrophenylphosphatase), AAEL007584 (*trunk*), AAEL005635 (*Nup50*), AAEL003877 (*polyubiquitin)*, and AAEL006511 (*ubiquitin L40),* due to their constitutive high levels of expression during many developmental life stages (AAEL003877, AAEL006511, AAEL005635), or high levels of expression in the ovary triggered by uptake of a blood meal (AAEL010097, AAEL007097, AAEL007584), as evidenced from previous promoter characterization experiments or developmental transcriptional profiling (Figure 1A,B)(52–54). We inserted these various promoter fragments into a *piggybac* transposon upstream of the coding sequence for spCas9. Downstream to the promoter driven Cas9 we included a self-cleaving T2A peptide and eGFP coding sequence, together serving as a visual indicator of promoter activity. We also included a baculovirus derived Opie2 promoter (55) driving dsRed expression serving as a transgenesis marker (Figure 1C). The *piggybac* transgenes were injected into the germline of wildtype embryos (*Ae. aegypti* genome sequence strain Liverpool) (56), and transgenic mosquitoes harboring these transgenes were readily identified by bright dsRed fluorescent expression in the abdomen (Supplementary Figure 1). We established stable transgenic strains and dissected the germline tissues from each strain to assess promoter activity and found moderate levels of eGFP present in the ovary of lines AAEL010097-Cas9, AAEL007584-Cas9, AAEL005635-Cas9 and AAEL006511-Cas9,. TheAAEL010097-Cas9 and AAEL006511-Cas9 lines also exhibited only weak expression in the testes (data not shown). Finally, the AAEL003877-Cas9 line exhibited no detectable eGFP signal, consistent with previous work indicating this promoter does not express in the germline (53) (Figure 1D). Importantly, we were able to establish homozygous stocks for each line demonstrating that no significant toxicity is associated with high transgenic expression of Cas9.

**Figure 1.**
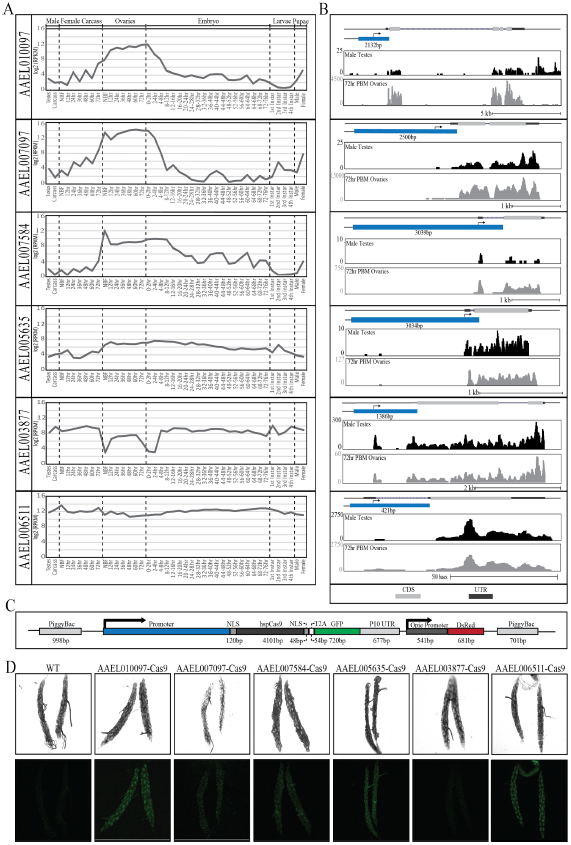
Rationally chosen, native promoters drive strong expression of Cas9. Log_2_ RPKM expression values for AAEL010097, AAEL007097, AAEL007584, AAEL005635, AAEL003877 and AAEL006511 were plotted across development. Samples include, from left to right: testis; male carcasses (lacking testes); carcasses of females prior to blood feeding (NBF), and at 12 hr, 24 hr, 36 hr, 48 hr, and 72 hr post-blood meal (PBM); ovaries from NBF females and at 12 hr, 24 hr, 36 hr, 48 hr, and 72 hr PBM; embryos from 0–2 hrs through 72–76 hr; whole larvae from 1st instar, 2nd instar, 3rd instar and 4th instar; male pupae; female pupae (A). Genome browser snapshots of AAEL010097, AAEL007097, AAEL007584, AAEL005635, AAEL003877 and AAEL006511, including expression tracks for both 72hr PBM ovaries and testes. The light gray box indicates coding sequence; blue box represents promoter element with length indicated in bp; Y axis shows the expression level based on raw read counts (B). Schematic representation of the piggybac-mediated Cas9 construct including the *Streptococcus pyogenes Cas9*-T2A-eGFP gene driven by selected promoters (blue), dsRed expressed under the control of the Opie2 promoter, which serves as a transgenesis marker. NLS represent nuclear localization signal (C). Confocal images using white light or GFP-flitering of transgenic Cas9 line ovaries (D).

### Germline expression of Cas9 in *Ae. aegypti* increases targeting efficiency

To test the efficacy of our transgenic *Ae. aegypti* Cas9 strains, we first attempted to mutate the *kynurenine hydroxylase* (*kh*) gene (AAEL008879), which was previously characterized as having an easily detectable dominant eye phenotype as the loss of *kh* gene function leads to severe eye pigmentation defects (31). To do so, we designed a single-guide RNA (Kh-sgRNA) targeting exon 4 of the *kh* gene, and to test its basic functionality we co-injected this *in vitro* transcribed Kh-sgRNA into 450 wildtype embryos (referred to as generation 0 [G_0_]) along with purified recombinant Cas9 protein. Following injection, complete white-eye and partial mosaic white-eye mutants were readily observed in adults (Supplementary Figure 2). The G_0_s had a survival rate of 35±7% and a mutagenesis efficiency of 17±3%, and a mutant heritability rate of 33±7% in the G_1_ generation (Table 1). Once we confirmed functionality of the Kh-sgRNA using purified recombinant Cas9 protein, we then injected this Kh-sgRNA separately into 450 embryos (three injections of 150 embryos in biological triplicate) collected from each of our transgenic *Ae. aegypti* Cas9 strains without including recombinant Cas9 protein in the injection mix. Complete white eye and mosaic white eye mutants were readily observed in injected G_0_ mosquitoes from 5/6 lines including AAEL010097-Cas9, AAEL007097-Cas9,AAEL007584-Cas9, AAEL005635-Cas9 and AAEL006511-Cas9. Notably, directly injecting Kh-sgRNAs into these transgenic Cas9 expression lines resulted in significantly higher G_0_ survival rates (61±7%, 53±4%, 63±4%, 64±6%, and 63±4%, respectively), increased G_0_mutagenesis efficiencies (85±5%, 27±6%, 52±7%, 47±3% and 66±4%, respectively), and increased overall heritable G_1_ mutation rates (60±9%, 59±6%, 59±5%, 60±6%, 66±3%, respectively) compared to when Kh-sgRNA was co-injected with recombinant Cas9 protein into wildtype embryos (Table 1). As reported previously, homozygous viable *kh* mutants have a dramatic eye pigmentation defect that can be visualized in larvae, pupae, and adults (Figure 2A)(31). To confirm the phenotypic defects described above were due to mutagenesis of the *kh* gene, genomic DNA spanning the target site was amplified from homozygous mutant G_1_s (Figure 2A) and sequenced, confirming the presence of insertion/deletions (indels) in the Kh-sgRNA genomic DNA target site (Supplementary Figure 3A,B). In contrast, Kh-sgRNA injected directly into the AAEL003877-Cas9 strain failed to show any visible mutant phenotypes in the G_0_ generation, presumably due to lack of germline expression by this promoter (53).

**Table 1.** Summary of the injection, survival and mutagenesis rates mediated by Kh-sgRNA,W1-sgRNA, and Y-sgRNA in wild type and Cas9 expressing lines.

| *Ae. aegypti* line | Injected component* | # Injected G<sub>0</sub>s | G<sub>0</sub> Adult Survivors (%) | G<sub>0</sub> mosaic (%) | G<sub>1</sub> mutant adult (%)<sup>&</sup> |
| --- | --- | --- | --- | --- | --- |
| Liverpool (wildtype) | Kh-sgRNA, W1-sgRNA, Y-sgRNA | 450,450,450 | 67±4,63±5,61±4 | 0,0,0 | 0,0,0 |
| Liverpool (wildtype) | Cas9 protein | 450,450,450 | 39±4,41±4,44±7 | 0,0,0 | 0,0,0 |
| Liverpool (wildtype) | Kh-sgRNA, W1-sgRNA, Y-sgRNA /Cas9 protein | 450,450,450 | 35±7,40±6,30±8 | 17±3,21±3,18±6 | 33±7,32±5,30±6 |
| AAEL010097-Cas9 | Kh-sgRNA, W1-sgRNA, Y-sgRNA | 450,450,450 | 61±7,71±6,65±7 | 85±5,87±6,90±5 | 60±9,69±4,62±4 |
| AAEL007097-Cas9 | Kh-sgRNA, W1-sgRNA, Y-sgRNA | 450,450,450 | 53±4,58±4,62±9 | 27±6,33±4,31±5 | 59±6,56±6,62±5 |
| AAEL007584-Cas9 | Kh-sgRNA, W1-sgRNA, Y-sgRNA | 450,450,450 | 63±4,55±7,61±5 | 52±7,63±8,57±5 | 59±5,51±7,57±9 |
| AAEL005635-Cas9 | Kh-sgRNA, W1-sgRNA, Y-sgRNA | 450,450,450 | 64±6,67±3,55±4 | 47±3,62±7,49±4 | 60±6,50±6,54±4 |
| AAEL003877-Cas9 | Kh-sgRNA, W1-sgRNA, Y-sgRNA | 450,450,450 | 51±9,61±8,61±7 | 0,0,0 | 0,0,0 |
| AAEL006511-Cas9 | Kh-sgRNA, W1-sgRNA, Y-sgRNA | 450,450,450 | 63±4,57±5,59±6 | 66±4,73±6,70±7 | 66±3,65±3,67±4 |

\*sgRNA 100 ng/ul; Cas9 protein 300 ng/ul.
<sup>&</sup> The overall heritable mutation rate was calculated as the number of mutant G<sub>1</sub>s divided by the number of all G<sub>1</sub>s observed.
Biological triplicate consisting of three independent injections (each injection replicate with 150 embryos total 450 embryos) were performed and the results are shown as the mean ± SE throughout table.

**Figure 2.**
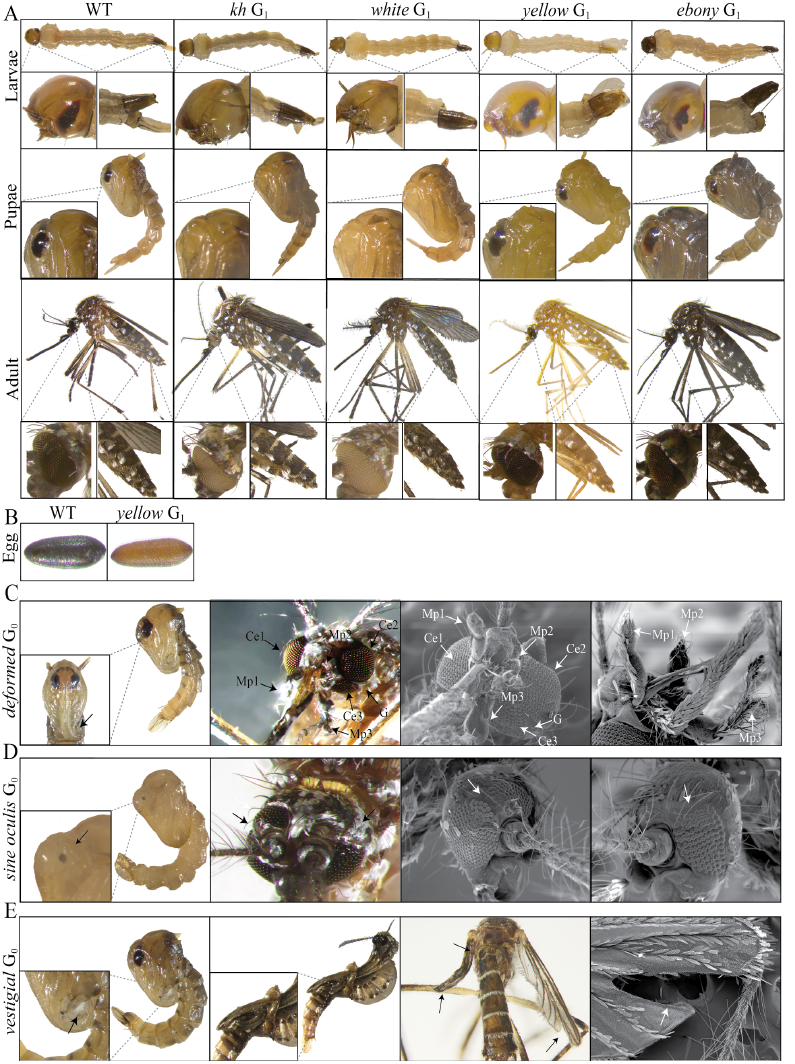
Severe mutant phenotypes caused by CRISPR/Cas9 mediated disruption. Larva, pupae, and adult phenotypes of wild type, *kynurenine hydroxylase (kh)*, *white*, *yellow* and *ebony* mutant G1’s, respectively, with clearly distinguishable eye (*kh* and *white*) and cuticle pigment (*yellow* and *ebony*) defects (A). Embryo phenotypes of wild type (left) and *yellow* mutants (right) (B). Pupae and adult scanning electron microscopy (SEM) images of the head of the *deformed* G0 mutants with three compound eyes (Ce), three maxillary palps (Mp), furrowed eyes and deformed mouthparts (arrows) (C). Pupae and adult SEM images of the head of the *sine oculis* G0 mutants. Arrows point to the ectopic eyes (D). Images of pupae and adult wings of *vestigial* G0 mutants. Arrows point to pronounced wing, halteres, and forked wing defects (E)

Given that CRISPR/Cas9 targeting efficiency varies significantly between loci and even between target sites within the same locus (26, 27, 30, 36), we wanted to further investigate the efficacy of these transgenic Cas9 expressing strains. We therefore designed two additional sgRNAs targeting uncharacterized conserved genes that may be useful for developing future control methodologies *Ae. aegypti.* One sgRNA (W1-sgRNA) was designed to target exon 3 of the *white* gene (AAEL016999) (Supplementary Figure 4A) which is the *Aedes* 1:1 orthologue to *D. melanogaster white* and functions as an ATP-binding cassette transporter important for red and brown eye color pigmentation (57, 58). Another sgRNA was designed to target exon 2 of the *Yellow* gene (AAEL006830) (Supplementary Figure 5A) which is the *Aedes* 1:1 orthologue to *D. melanogaster yellow*, a gene important for melanization of the cuticle (59, 60). To initially test their functionality, we co-injected either *in vitro* transcribed W-sgRNA or Y-sgRNA, separately, into 450 wildtype embryos (three injections of 150 embryos in biological triplicate) each along with purified recombinant Cas9 protein. Following injection, complete white eye and partial mosaic white eye, or complete yellow body and partial mosaic yellow body mutants were readily observed in G_0_s indicating that the sgRNAs are functional (Table 1, Supplementary Figure 2A). Once sgRNA functionality was confirmed using recombinant Cas9 protein, we then injected these sgRNAs separately into 450 embryos (three injections of 150 embryos in biological triplicate) collected from each of our transgenic *Ae. aegypti* Cas9 expressing strains without including recombinant Cas9 protein in the injection mix. Similar to the Kh-sgRNA results described above, remarkably higher survival rates, mutagenesis efficiencies, and heritable mutation rates were also achieved for both W-sgRNA and Y-sgRNA when injected into all transgenic Cas9 lines, except AAEL003877-Cas9 (Table 1). As in other Dipterans, homozygous G_1_ mutants for *white* have dramatic eye pigmentation defects, similar to the *kh* mutants, that can be visualized in larvae, pupa, and adults (Figure 2A). Additionally, homozygous G_1_ mutants for *yellow* have a striking yellow cuticle pigmentation defect that can be visualized in, larvae, pupae, adults, and even eggs (Figure 2A,B). To confirm the phenotypic defects from these sgRNAs were due to mutagenesis of the *white* and *yellow* genes, genomic DNA from homozygous mutant G_1_s (Figure 2A) was amplified spanning the target sites and sequenced confirming the presence of insertion/deletions (indels) in both the W-sgRNA and Y-sgRNA genomic DNA target sites (Supplementary Figure 4A,B, 5A,B). Altogether, these results indicate that the *Ae. aegypti* transgenic Cas9 expression system that we generated can effectively express Cas9 protein, and direct injection of *in vitro* transcribed sgRNAs is sufficient to rapidly disrupt function of target genes of interest and generate highly heritable mutations. Out of all the transgenic lines, the AAEL010097-Cas9 line consistently showed the highest survival rate and efficiency of mutagenesis. Therefore, we performed all subsequent experiments using this strain.

### Precise gene disruption in *Ae. aegypti*

To further measure the efficacy of the AAEL010097-Cas9 line, we decided to target additional genes causing readily visible phenotypes in related organisms that have yet to be studied in *Ae. aegypti*. Specifically, we chose genes AAEL005793, AAEL009950, AAEL009170 and AAEL003240, which are conserved 1:1 orthologues with *the D. melanogaster* genes *ebony*, *deformed*, *vestigial* and *sine oculis*, respectively. *Ebony* encodes the enzyme N-β-alanyl dopamine (NBAD) synthetase that converts dopamine to NBAD. Loss of ebony function increases black cuticle pigment in *D. melanogaster* (61). *Deformed* is a homeobox-containing (Hox) transcription factor, loss of function of this gene results in dramatic defects in derivatives of the maxillary segments, mandibular segments, and anterior segments in *D. melanogaster* (62). *Vestigial* encodes a nuclear protein that plays a central role in the development of the wings and loss of *vestigial* results in the failure of proper wing development in *D. melanogaster* (63, 64). *Sine oculis* locus encodes a homeodomain containing protein, which plays an essential role in eye development and loss of *sine oculis* function leads disruption of the eye disc formation in *D. melanogaster* (65, 66).

To disrupt these gene, a single sgRNA was designed (E-sgRNA, D-sgRNA, So-sgRNA, V-sgRNA) to target an exon in each of these genes (Supplementary Figures 6-9), and then individually injected into 200 AEL010097-Cas9 embryos. Injection of E-sgRNA resulted in 82% of G_0_s showing pronounced dark coloration of the adult cuticle (Supplementary Figure 2). These mutations were heritable and we established homozygous viable stocks by pairwise mating G_1_s (Table 2, Figure 2A). D-sgRNA injection resulted in the death of more than 72% of G_0_ mosquitoes prior to the pupal stage indicating that this gene is critical for proper development.However, 87% of the mosquitoes that reached adulthood exhibited severe mutant phenotypes including an extra eye (triple eyes), mal-shaped proboscis, and even an extra maxillary palp (triple maxillary palps) (Table 2, Figure 2C). Unsurprisingly, all surviving G_0_ mutant females failed to blood feed, preventing us from establishing homozygous mutant lines. In So-sgRNA injected mosquitoes, 61% of the surviving G_0_s had clearly visible eye defects do to cell death in the primordium region (Table 2, Figure 2D), As with the *deformed* mutants, we were unable to establish homozygous *sine oculis* mutant lines because these mutations were homozygous lethal. Finally, 70% of V-sgRNA-injected G_0_s died at the pupal stage, with many of the dead pupae showing deformed rudimentary wing appendages (Figure 2E). Of the few individuals that survived to adulthood, 75% had undeveloped wings and/or halteres (Table 2, Figure 2E), inhibiting their ability to fly or mate making it impossible to establish homozygous mutant lines for this gene. To confirm that the phenotypes described above were caused by disruption of the specific genes targeted by the sgRNAs (E-sgRNA, D-sgRNA, So-sgRNA, V-sgRNA), genomic DNA from the mutants (Figure 2C-E) was amplified spanning the target sites and sequenced confirming a selection of insertion/deletions (indels) in both the genomic DNA target sites (Supplementary Figures 6-9).

**Table 2.** Summary of the injection, survival and mutagenesis rates mediated by E-sgRNA,D-sgRNA, SO-sgRNA and V-sgRNA in AAEL010097-Cas9 line.

| *Ae. aegypti* line | Injected component* | # Injected G<sub>0</sub>s | G<sub>0</sub> larva Survivors (%) | G<sub>0</sub> Adult Survivors (%) | G<sub>0</sub> mutant adult (%) | G<sub>1</sub>s mutant adult (%)<sup>&</sup> |
| --- | --- | --- | --- | --- | --- | --- |
| AAEL010097-Cas9 | E-sgRNA | 200 | 115 (58) | 112 (56) | 92 (82) | 751 (67) |
| AAEL010097-Cas9 | D-sgRNA | 200 | 122 (61) | 33 (28) | 29 (87) | lethal |
| AAEL010097-Cas9 | So-sgRNA | 200 | 104 (52) | 98 (52) | 59 (61) | lethal |
| AAEL010097-Cas9 | V-sgRNA | 200 | 134 (67) | 40 (30) | 22 (75) | lethal |

\*sgRNA 100 ng/ul
<sup>&</sup> The overall heritable mutation rate was calculated as the number of mutant G<sub>1</sub>s divided by the number of all G<sub>1</sub>s observed.

### Multiple sgRNAs dramatically improve mutagenesis rates

To determine if simultaneous injection of two sgRNAs into the embryos of the AAEL010097-Cas9 line could induce higher mutagenesis efficiencies, and germline transmission rates, we designed another sgRNA (W2-sgRNA) also targeting exon 3 of the *white* gene (Supplementary Figure 10). Initially we tested functionality of the W2-sgRNA by co-injecting this guide with recombinant Cas9 protein into wildtype embryos. We found that this W2-sgRNA is efficient at generating mutations as 41±7% of injected embryos survived to adult, 19±6% of adult survivors showed the mosaic and white eye mutant phenotypes, and 27±5% G_1_s showed the mutant white eye phenotype (Table 3). Similar to results described above, directly injecting the W2-sgRNA into the embryos of the AAEL010097-Cas9 line resulted in higher survival rates (62±6%), greater mutagenesis efficiencies (67±9%), and higher germline transmission rates (56±9%) (Table 3). Importantly, when we co-injected W1-sgRNA (described above) and W2-sgRNA together into embryos of the AAEL010097-Cas9 line, *white* mutants were obtained in the G_0_s at an extremely higher frequency of 94±3% with a remarkable G_1_ transmission frequency of 96±3% (Table 3). Furthermore, genomic sequencing confirmed that some of the injections generated large deletions spanning the genomic distance (~350 bp) between the W1-sgRNA and W2-sgRNA target sites (Supplementary Figure 10). Together, these results indicate that by simultaneous injection of multiple sgRNAs targeting the same gene higher mutagenesis rates and large deletions can be readily achieved.

**Table 3.** Summary of the injection, survival and mutagenesis rates mediated by W1-sgRNA and W2-sgRNA in wild type and AAEL010097-Cas9 line

| *Ae. aegypti* line | Injected component* | # Injected G<sub>0</sub>s | G<sub>0</sub> Adult Survivors (%) | G<sub>0</sub> mosaic (%) | G<sub>1</sub>s mutant adult (%)<sup>&</sup> |
| --- | --- | --- | --- | --- | --- |
| Liverpool (wildtype) | W2-sgRNA /Cas9 protein | 450 | 41±7 | 19±6 | 27±5 |
| Liverpool (wildtype) | W1-sgRNA/W2-sgRNA /Cas9 protein | 450 | 37±5 | 34±5 | 37±9 |
| AAEL010097-Cas9 | W2-sgRNA | 450 | 62±6 | 67±9 | 56±9 |
| AAEL010097-Cas9 | W1-sgRNA/W2-sgRNA | 450 | 61±8 | 94±3 | 96±3 |

\*sgRNA 100 ng/ul; Cas9 protein 300 ng/ul.
<sup>&</sup> The overall heritable mutation rate was calculated as the number of mutant G<sub>1</sub>s divided by the number of all G<sub>1</sub>s observed.
Biological triplicate consisting of three independent injections (each injection replicate with 150 embryos total 450 embryos) were performed and the results are shown as the mean ± SE throughout table.

### Highly efficient multiplex gene disruption from a single injection

Given the laborious and time consuming crossing required to generate *Ae. aegypti* lines with mutations at multiple genes, we explored whether double or triple-gene knockout strains could quickly be generated in one step by co-injecting a multiplexed combination of sgRNAs targeting different genes. To do so, we made four mixes including three combinations of two sgRNAs including; W1-sgRNA and Y-sgRNA (mix 1), W1-sgRNA and E-sgRNA (mix 2), Y-sgRNA and E-sgRNA (mix 3), and one combination of three sgRNAs; W1-sgRNA, Y-sgRNA and E-sgRNA (mix 4). We injected these four mixes separately into 200 embryos from the AAEL010097-Cas9 line. We found that 90%, 87%, 93% and 94% of the G_0_ injected survivors contained at least one mutant phenotype for the injection sets 1-4, respectively (Table 4).Furthermore, 90%, 88% and 92% of the G_0_ injected survivors had double mutants when injected with sets 1-3, respectively (Table 4, Supplementary Figure 11). When we co-injected a combination of three sgRNA (mix 4), we found that 67% of the surviving G0 adults contained mutations in all three genes targeted (Table 4, Supplementary Figure 11). Furthermore, these mutations were heritable as homozygous viable mutant stocks following pairwise mating of G_1_ multi-mutants (Figure 3). Together these results demonstrate that simultaneous and heritable multiple gene disruptions can be efficiently achieved by using transgenic expression of Cas9 and this simple technique should facilitate the rapid understanding of gene function and help tease apart gene networks in this non-model organism.

**Table 4.** Single injection of multiple sgRNAs into the embryos of AAEL010097-Cas9 line results in highly efficient rates of multi-gene disruption.

| Injected sgRNAs | Inject ed embryos | G<sub>0</sub> Adult Survivors (%) | G<sub>0</sub> mosaic (%) | Single mutant G<sub>0</sub>s (%); G<sub>1</sub>s (%)<sup>#</sup> | | | Double mutant G<sub>0</sub>s (%); G<sub>1</sub>s (%)<sup>\$</sup> | | | Triple mutant G<sub>0</sub>s(%); G<sub>1</sub>s (%)<sup>*</sup> |
| --- | --- | --- | --- | --- | --- | --- | --- | --- | --- | --- |
|  |  |  |  | *W* | *Y* | *E* | *W/Y* | *W/E* | *Y/E* | *W/Y/E* |
| W1-sgRNA/Y-sgRNA | 200 | 122(61) | 110 (90) | 4(4); 170 (15) | 7(6); 238 (21) | N/A; N/A | 99(90); 488(43) | N/A; N/A | N/A; N/A | N/A; N/A |
| W1-sgRNA/E-sgRNA | 200 | 113(56) | 98(87) | 7(7); 141 (17) | N/A; N/A | 5(5); 224(27 ) | N/A; N/A | 86(88); 390 (47) | N/A; N/A | N/A; N/A |
| Y-sgRNA/E-sgRNA | 200 | 102(51) | 95(93) | N/A; N/A | 3(3); 343(25) | 5(5); 261 (15) | N/A; N/A | N/A; N/A | 87(92); 536 (39) | N/A; N/A |
| W1-sgRNA/Y-sgRNA/E-sgRNA | 200 | 86(43) | 81 (94) | 3(4); 85(11) | 3(4); 54(7) | 1(1); 23(3) | 5(6);131 (17) | 9(11);101(13) | 6(7); 162(21) | 54(67); 85(11) |

<sup>#</sup> Single mutant phenotype, *W*: white eye; *Y*: yellow body; *E*: dark body
<sup>\$</sup> Double mutant phenotypes, *W/Y*: white eye and yellow body, *W/E*: white eye and dark body; *Y/E* yellow body and dark body
<sup>\*</sup> Triple mutant phenotypes, *W/Y/E*: white eye, yellow body, and dark body

**Figure 3.**
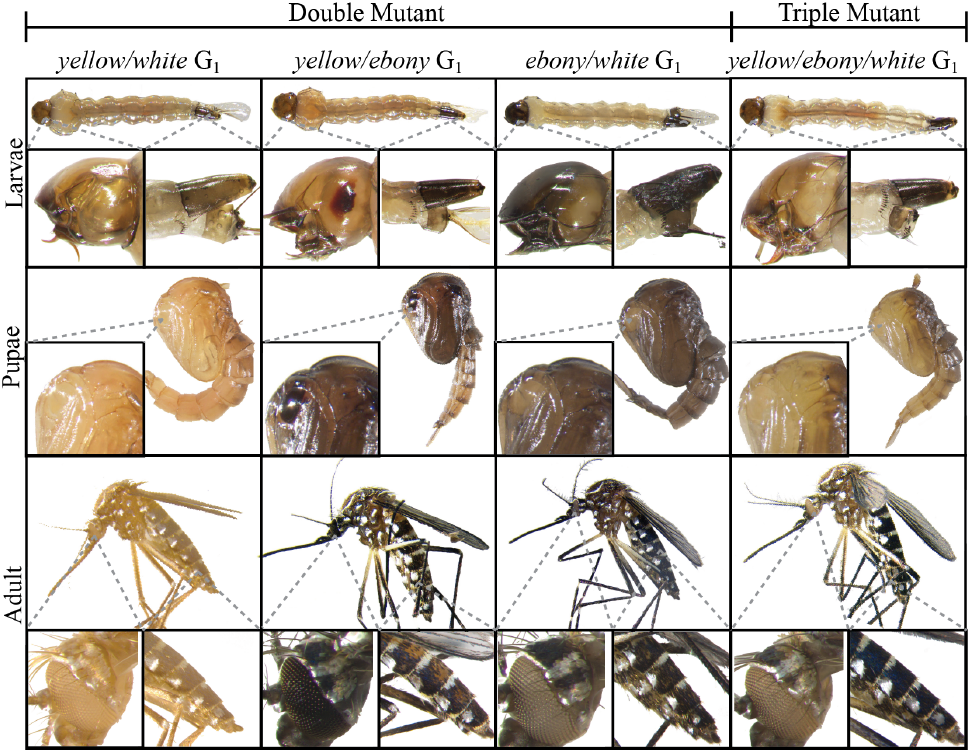
Single injections of multiplexed sgRNAa robustly generate double and triple mutant mosquitoes. Larva, pupae, and adult G1 phenotypes for double mutants including: yellow body and white eyes (*yellow/white)*, a mixture of yellow and dark body (*yellow/ebony)*, dark body and white eyes (*ebony/white)*, and one triple mutant which is a phenotypic mixture of yellow and dark body and white eyes (*yellow/ebony/white)*. The striking differences between wild type and mutant larva, pupae and adult are highlighted.

### Increased rates of HDR with dsDNA donors

Given the exceedingly high rate of NHEJ-induced mutation when using theAAEL010097-Cas9 line, we wanted to assess knock-in rates mediated by HDR via dsDNA donors in this line. Our strategy was to employ two donor plasmids, a *white*-donor plasmid based on the W1-sgRNA and a *kh*-donor plasmid based on the Kh-sgRNA. Each of these two donor plasmids was designed to contain a dominant fluorescent marker consisting of 3xP3-dsRed, expressing in the larvae and adult photoreceptors, flanked by ~1kb homology arms that were derived from the genomic sequence immediately flanking the target cleavage sites (Figure 4A,B). We directly compared two approaches; i) co-injection of dsDNA donor combined with *in vitro* transcribed sgRNA and purified recombinant Cas9 protein into wildtype embryos; and ii) co-injection of circular dsDNA donor combined with *in vitro* transcribed sgRNA injected into AAEL010097-Cas9 line embryos. For approach i, a total of 600 embryos were separately injected for both the *white-*donor and *kh*-donor, with a HDR rate of 0.15% and 0.14% (Table 5), respectively. For approach ii, a total of 600 embryos from the *Ae. aegypti* AAEL010097-Cas9 line were separately injected for both the *white-*donor and *kh*-donor, with a HDR rate of 2.36% and 2.48% (Table 5), respectively. Overall, we see a dramatic, yet consistent, 15-17 fold increase in rates of HDR when using the AAEL010097-Cas9 line compared to the non-transgenic method of supplying Cas9. Gene-specific insertion of our donor cassettes into the intended target sites was confirmed by subsequent genomic PCR and sequencing (Figure 4A-C, Supplementary Figure 12).

**Table 5.** Transgenic source of Cas9 results in increased rates of HDR with dsDNA donors.

| *Ae. aegypti* strain | Injected component* | Injected embryos | Adult G<sub>0</sub> Survivors (%) | HDR event<sup>s</sup> |  |
| --- | --- | --- | --- | --- | --- |
|  |  |  |  | # group founders/total group (%) | # HDR G1/total G1 (%) |
| **Liverpool (wildtype)** | W1-sgRNA/Cas9/*white*-donor | 600 | 251(41.83) | 1/20 (5) | 9/5371 (0.15) |
| **Liverpool (wildtype)** | Kh-sgRNA/Cas9/*kh*-donor | 600 | 273 (45.50) | 1/20 (5) | 7/4773 (0.14) |
| **AAEL010097-Cas9** | W1-sgRNA/*white*-donor | 600 | 308 (51.33) | 4/20 (20) | 118/4993 (2.36) |
| **AAEL010097-Cas9** | Kh-sgRNA/*kh*-donor | 600 | 298 (49.67) | 5/20 (25) | 149/6017 (2.48) |

\* The concentration of sgRNA is 100 ng/ul; Cas9 protein 300 ng/ul. *white*-donor and *kh*-donor plasmids 100 ng/ul
<sup>s</sup> HDR represent homology-directed repair

**Figure 4.**
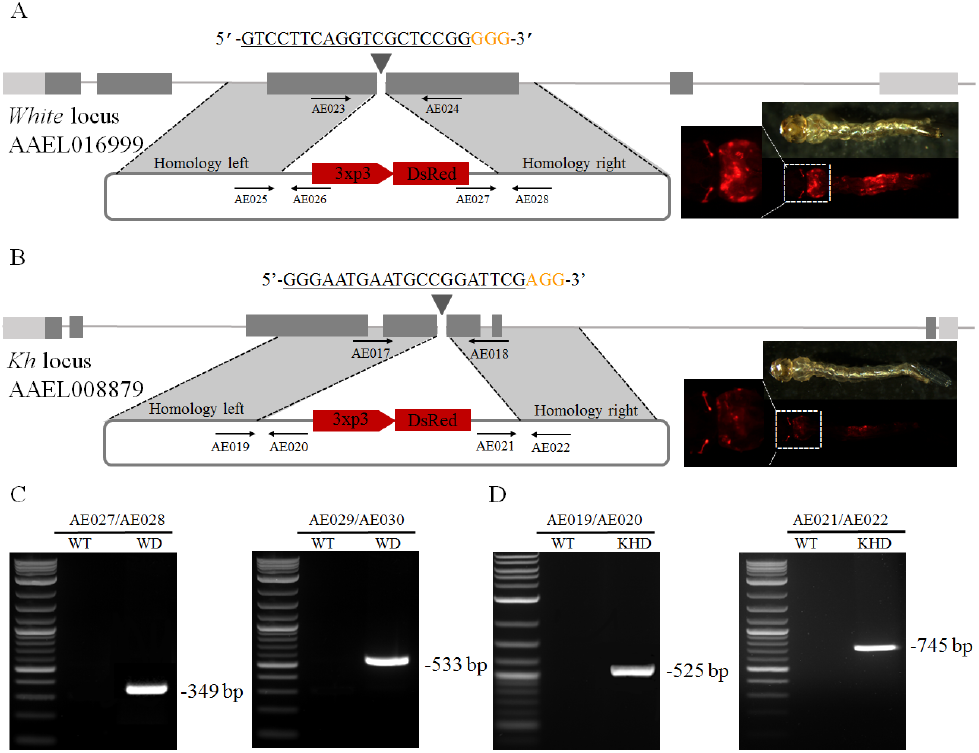
Highly efficient site-specific integration via CRISPR mediated HDR. Schematic representations of the *white* locus and *white*-donor construct (A), and the *kynurenine hydroxylase* locus and *kh*-donor construct (B). Exons are shown as boxes, coding regions are depicted in black and the 5’and 3’ UTRs in gray. Locations and sequences of the sgRNA targets are indicated with the PAM shown in yellow. Black arrows indicate approximate positions and directions of the oligonucleotide primers used in the study. The donor plasmids (blue) express fluorescent eye marker (3xp3-DsRed) inserted between regions of homology from the *white* and *kynurenine hydroxylase* locus, respectively (A,B). Gene amplification analysis confirms site-specific integration of the *white*-donor construct into the *white* locus using combinations of genomic and plasmid donor specific primers (933Cms3/933Cms4 expected 349bp, and 933Cms5/933Cms6 expected 533bp) (C), and also confirms the integration of the *kh*-donor construct into the *kh* locus using combinations of genomic and plasmid donor specific primers (924ms3/924ms4 expected 525bp, and 924ms5/924ms6 expected 745bp) with no amplification in wildtype (D). WT represents wild-type, KD represents knockin with white-donor, KHD represents knockin with *kh-donor.*

## Discussion

Previous studies have demonstrated effective CRISPR/Cas9 genome editing in the mosquito *Ae. aegypti (30–34)*, however these studies utilized a non-transgenic source of Cas9 limiting both survivorship and editing efficiencies. To overcome these previous limitations, and to reduce the complexity of injecting multiple components (i.e. Cas9 and sgRNA), here we developed a simplified transgenic Cas9 expression system in *Ae. aegypti,* similarly to what is routinely used in other organisms such as *D. melanogaster (27, 41, 42)*. Importantly, to achieve highly specific and consistent genome modifications, we demonstrate that embryos from these Cas9 strains need only be injected with easy-to-make sgRNAs. By using these Cas9 strains, we disrupted multiple genes that were either homozygous viable (*kh*, *white*, *yellow*, and *ebony*), or homozygous lethal (*deformed*, *sine oculis*, *vestigial)* resulting in dramatic phenotypes affecting viability, vision, flight, and blood feeding, and therefore may be useful for developing novel control strategies and genetic sexing techniques in the future. For example, in a *yellow* mutant background, the endogenous gene encoding *yellow* could be linked to the male determining locus (32) using CRISPR-mediated HDR, to generate a robust genetic sexing system by which male embryos/larvae/adults would be dark and female embryos/larvae/adults would be yellow.

An appealing advantage of our Cas9 transgenic system is the ability to efficiently disrupt multiple genes simultaneously. We have demonstrated that we can efficiently generate large deletions, or even double (*yellow-white; ebony-white; yellow-ebony*), or triple (*yellow-ebony-white*) mutants. Importantly, these multi-mutants can rapidly be generated in a single-step approach by injecting multiple sgRNAs into the embryos of the transgenic Cas9 strains, significantly reducing downstream efforts. This rapid multiplex gene knockout approach will be instrumental for dissecting gene networks in this non-model organism.

The germline Cas9 strains developed here may also bring us one step closer to engineering an effective CRISPR/Cas9-homing based gene drive system (47, 48) in this organism. Homing based drive systems rely on HDR to convert heterozygous alleles into homozygous alleles in the germline, and have recently been successfully engineered in two *Anopheline* mosquito species (51, 67). While these studies were fruitful at significantly biasing rates of Mendelian inheritance rates of the drive containing alleles, they were severely limited by the rapid evolution of resistance alleles generated by NHEJ, and are therefore not predicted to spread into diverse wild populations (68). It was hypothesized that these resistance alleles formed due to high levels of maternal deposition of Cas9 in the embryo, and by restricting Cas9 expression to the germline may subsequently increase rates of HDR and reduce rates of NHEJ. In addition to restricting expression to the germline, multiplexing of sgRNAs in the drive, and designing the drive to target a critical gene, have also been proposed as innovative strategies to increase rates of HDR and reduce resistance caused by NHEJ (47, 48, 68), however these hypotheses remain to be demonstrated. Notwithstanding, it will be interesting to determine if our *Ae. aegypti* Cas9 strains, each with varying expression in the germline, will be effective in a gene drive system designed for *Ae. aegypti*. This would be straightforward to test in a molecularly confined safe split-gene drive design where the Cas9 and the drive are positioned at different genomic loci. In this split-design, the Cas9 strains we developed can be directly tested without further modification by simply genetically crossing with a split gRNA-drive component and measuring rates of inheritance (49, 69).

While the CRISPR/Cas9 transgenic system developed here is quite effective, it would be useful to have the ability to supply the sgRNAs transgenically. In *D. melanogaster,* polymerase-3 promoters have been utilized to express sgRNAs, and through genetic crosses with Cas9 strains mutation efficiency could be increased up to 100% (42, 43). Therefore, we intend to focus future efforts on developing transgenic methods to supply sgRNAs in an effort to further improve mutagenesis efficiencies and also to develop effective gene drives in the organism. Overall, our results demonstrate that our simplified transgenic Cas9 system has improved capacity to rapidly induce highly efficient and specific targeted genome modifications including gene disruptions, deletions, and insertions. Given their high efficiencies, these Cas9 strains can be used to quickly generate novel genome modifications allowing for high-throughput gene targeting, thereby accelerating comprehensive functional annotation of the *Ae. aegypti* genome.

## Materials and methods

### Insect rearing

Mosquitoes used in all experiments were derived from of the *Ae. aegypti* Liverpool strain, which was the source strain for the reference genome sequence (56). Mosquitoes were raised in incubators at 28°C with 70–80% humidity and a 12 hour light/dark cycle. Larvae were fed ground fish food (TetraMin Tropical Flakes, Tetra Werke, Melle, Germany) and adults were fed with 0.3M aqueous sucrose. Adult females were blood fed three to five days after eclosion using anesthetized mice. All animals were handled in accordance with the guide for the care and use of laboratory animals as recommended by the National Institutes of Health and supervised by the local Institutional Animal Care and Use Committee.

### *piggyBac* mediated Cas9 plasmid construction

To generate the Cas9 constructs, the *piggyBac* plasmid pBac-3xP3-dsRed (a kind gift from R. Harrell) was digested and linearized with Fse1/Asc1 and this was used as a backbone for construct assembly using the Gibson method (70). The Opie2 promoter region (55) was amplified from vector pIZ/V5-His/CAT (Invitrogen) using primers AE001 and AE002, to drive the expression of dsRed. The promoters that we selected in the current study were based on three parameters: 1) an optimal location for primer design for ideal PCR conditions, 2) include as much regulatory information as possible without venturing into other genes or transcribed features as identified through (www.vector.caltech.edu), and 3) limit the size of our transgene to ensure integration events would occur at high frequency. A total of 2133 bp putative promoter region for gene AAEL010097, a 2500 bp putative promoter region for gene AAEL007097, a 3041 bp putative promoter region for gene AAEL007584, a 3034 bp putative promoter region for gene AAEL005635, a 1386 bp putative promoter region for gene AAEL003877, a 421 bp putative promoter region for gene AAEL006511 was PCR amplified from *Ae. aegypti* genomic DNA with primers AE003 and AE004, AE005 and AE006, AE007 and AE008, AE009 and AE010, AE011 and AE012, AE013 and AE014, respectively (52, 53). Downstream of each promoter element, a NLS-Cas9-NLS-T2A-eGFP-p10 UTR which consisted of elements i) 120 bp N-terminal nuclear localization signal (NLS) sequence; ii) a 4101 bp human codon optimized Cas9 (hspCas9) in which the sequence was derived from plasmid p3xP3-EGFP/*vasa*-3xFLAG-NLS-Cas9-NLS (44); iii) a 48 bp C-NLS, a 54 bp self cleaving peptide 2A (T2A); iv) a 720 bp green fluorescent protein (eGFP). A template DNA containing elements i-vi was generated using gene synthesis (Genescript), and subcloned into plasmids to be driven by promoters mentioned above. A 677 bp p10 3’ untranslated region (UTR) (71) was amplified with primers AE015 and AE016 from vector pJFRC81-10XUAS-IVS-Syn21-GFP-p10 and included as the 3’ UTR for NLS-Cas9-NLS-T2A-eGFP(Addgene plasmid 36432). All plasmids were grown in strain JM109 chemically competent cells (Zymo Research #T3005), and isolated using Zyppy Plasmid Miniprep (Zymo Research #D4037) and Maxiprep (Zymo Research #D4028) kits; the full plasmid sequence was verified using Source Bioscience Sanger sequencing services. A list of primer sequences used in the above construct assembly can be found in Supplementary Table 1.

### Generation of *Ae. aegypti* Cas9 transgenic lines

Transgenic *Ae. aegypti* Cas9 mosquitoes were created by injecting 0-1 hr old pre-blastoderm stage embryos with a mixture of *piggybac* vector containing the Cas9 expressing plasmid designed above (200ng/ul) and a source of *piggyBac* transposase (phsp-Pbac, (200ng/ul)) (72–74). Embryonic collection and microinjections were largely performed following previously established procedures (75). The injected embryos were hatched in deoxygenated H_2_0, and the surviving adults were put into cages, and blood feed 4 days later after eclosion. The larvae with positive fluorescent signals were selected under the fluorescent stereo microscope (Leica M165FC) for at least 8 generations to establish stable homozygous line.

### Production of sgRNAs

Linear double-stranded DNA templates for all sgRNAs were generated by template-free PCR with NEB Q5 high-fidelity DNA polymerase (catalog # M0491S) by combining primer pairs (sgRNA F and sgRNA R) following (29, 30). PCR reactions were heated to 98°C for 30 seconds, followed by 35 cycles of 98°C for 10 seconds, 58°C for 10 seconds, and 72°C for 10 seconds, then 72°C for 2 minutes. PCR products were purified with Beckman Coulter Ampure XP beads (catalog #A63880). Following PCR, sgRNAs were synthesized using the Ambion Megascript T7 in vitro transcription kit (catalog # AM1334, Life Technologies) according to the manufacturer’s protocols using 300ng of purified DNA template overnight at 37°C. Following in vitro transcription, the sgRNAs were purified with MegaClear Kit (catalog #AM1908, Life Technologies) and diluted to 1000 ng/ul in nuclease-free water and stored in aliquots at −80°C. Recombinant Cas9 protein from *Streptococcus pyogenes* was obtained commercially (CP01, PNA Bio Inc) and diluted to 1000 ng/ul in nuclease-free water and stored in aliquots at −80°C. All primer sequences can be found in table S1.

### Donor plasmids construction

We designed two donor plasmids, *white*-donor and *kh*-donor based on the sgRNAs (W1-sgRNA and Kh-sgRNA) that target the *white* and *kynurenine hydroxylase*c (*kh*) gene, respectively.Donor plasmids were constructed using standard molecular biology techniques based on Gibson assembly (70). The plasmids contained the following elements: (1) a 3xP3-DsRed fragment, which was amplified from vector pBac-3xP3-dsRed using primers AE039 and AE040 and expresses the dsRed dominant fluorescence marker visible in larvae photoreceptors as well as adult eyes, (2) DNA fragments ~ 1 kb in length each that are homologous to the *Ae. aegypti white* and *kh* locus immediately adjacent to the 5’ and 3’ ends of the W1-sgRNA and Kh-sgRNA target cutting site, amplified from *Ae. aegypti* genome DNA with primer pairs of AE041 and AE042, AE043 and AE044, AE045 and AE046, AE047 and AE048, respectively.

### CRISPR mediated microinjections

Embryonic collection and CRISPR microinjections were performed following previously established procedures (30, 75). The concentration of components used in the study was as follows; Cas9 protein at 300 ng/ul, sgRNA at 100 ng/ul, and donor plasmid at 100 ng/ul. To identify mutants, injected G_0_s and G_1_s were visualized at the life stages of larva, pupae, and adult carefully under a dissecting microscope (Olympus, SZ51). The heritable mutation rates were calculated as the number of mutant G_1_s out of the number of all G_1_ progeny crossed with respective mutant stocks. To molecularly characterize the induced mutations, genomic DNA was extracted from an individual mosquito with the DNeasy blood & tissue kit (QIAGEN) following the manufacturer's protocol. Target loci were amplified by PCR, PCR products were gel purified and were either sent directly for Sanger sequencing or cloned (TOPO-TA, Invitrogen). In the latter case, single colonies were selected and plasmids were extracted (Zyppy^™^ plasmid miniprep kit) and sequenced. Mutated alleles were identified by comparison with the wild-type sequence. Primers used for PCR and sequencing are listed in supplementary table 1. All photographs were obtained using fluorescent stereo microscope (Leica M165FC) and confocal microscope (Leica SP5).

## Author Contribution(s)

M.L. and O.S.A developed the protocol and designed experiments. M.L., M.B, T.Y. performed all experiments. M.L., M.B., B.J.W., and O.S.A contributed to the writing of the manuscript. All authors analyzed the data and approved final manuscript.

### Acknowledgements

This work was supported by a private donation from www.MaxMind.com to O.S.A., US National Institutes of Health (NIH) K22 grant to O.S.A. (5K22AI113060), a NIH R21 grant (1R21AI123937) to OSA, a NIH R21 grant (1R21AI115271) to B.J.W., and a DARPA Safe Genes Program Grant (HR0011-17-2-0047) to O.S.A. We are thankful to Robert Harrell for providing the pBac-3xP3-dsRed and phsp-Pbac plasmids, and Kate M. O’Connor-Giles for providing the p3xP3-EGFP/*vasa*-3xFLAG-NLS-Cas9-NLS plasmid and sequence, and to Anthony A. James for reading over the manuscript and providing insightful comments and edits.

## Disclosure

The authors declare no competing financial interests.

## All Figures and Tables

**Supplementary Figure 1.**
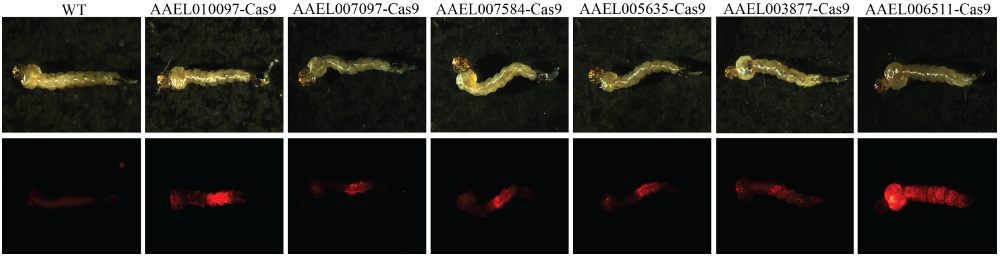
Cas9 expressing transgenic mosquitoes with dsRed selectable marker. Transgenic mosquitoes for each Cas9 strain showing the Opie2-dsRed fluorescent marker. Brightfield images (top) and corresponding dsRed fluorescent images showing robust expression in the larval gut and cuticle tissues.

**Supplementary Figure 2.**
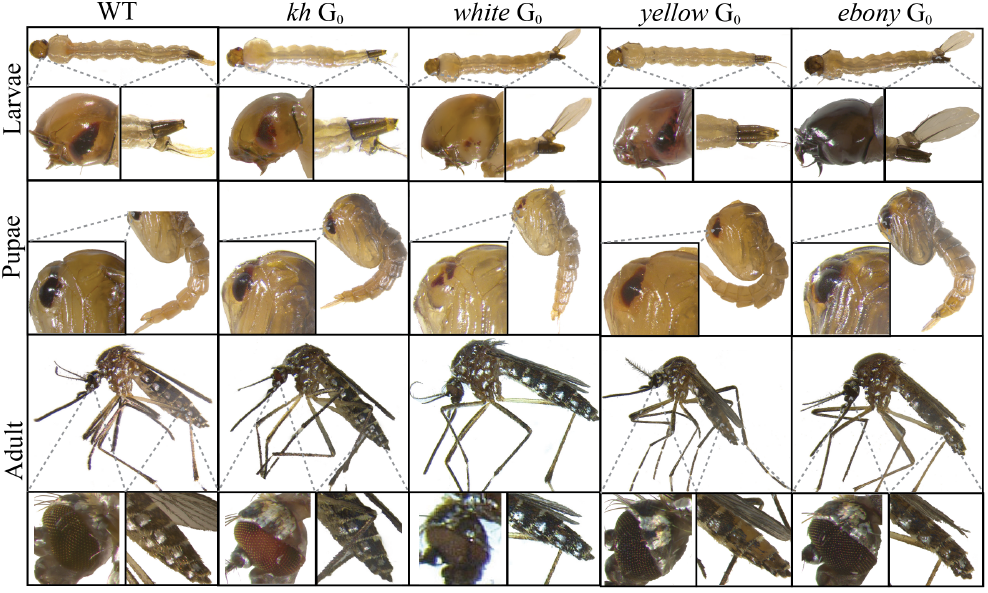
CRISPR/Cas9 induced G0 mosaic mutant phenotypes from single injections. Larval, pupae, and adult G0 mosaic phenotypes of wild type, *kh*, *white*, *yellow* and *ebony* mutants.

**Supplementary Figure 3.**
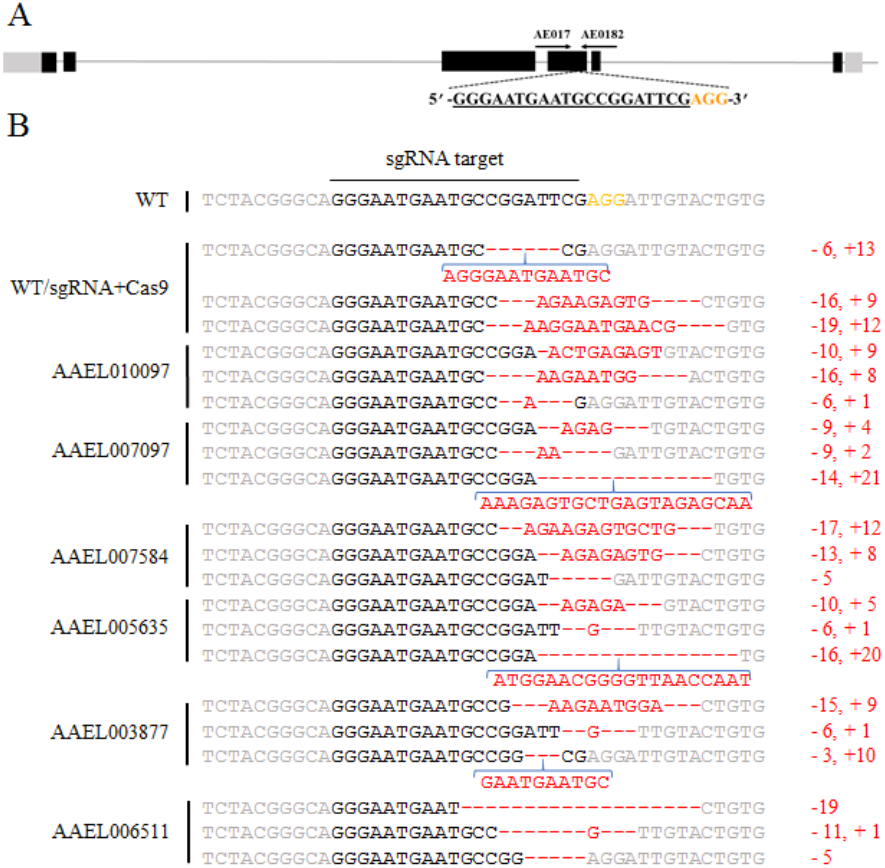
Confirmed mutagenesis of the *kh* locus. Schematic representation of the *kh* locus with exons indicated as boxes, coding regions depicted in black, and the 5’and 3’UTR’s in gray. Locations and sequences of the sgRNA targets are indicated with the PAM highlighted in yellow (A). Genomic sequencing analysis of indels from individuals sequenced from the various Cas9 strain injections. Top line represents WT sequence; PAM sequences (NGG) are indicated in yellow, and *kh* gene disruptions resulting from insertions/deletions are indicated in red (B).

**Supplementary Figure 4.**
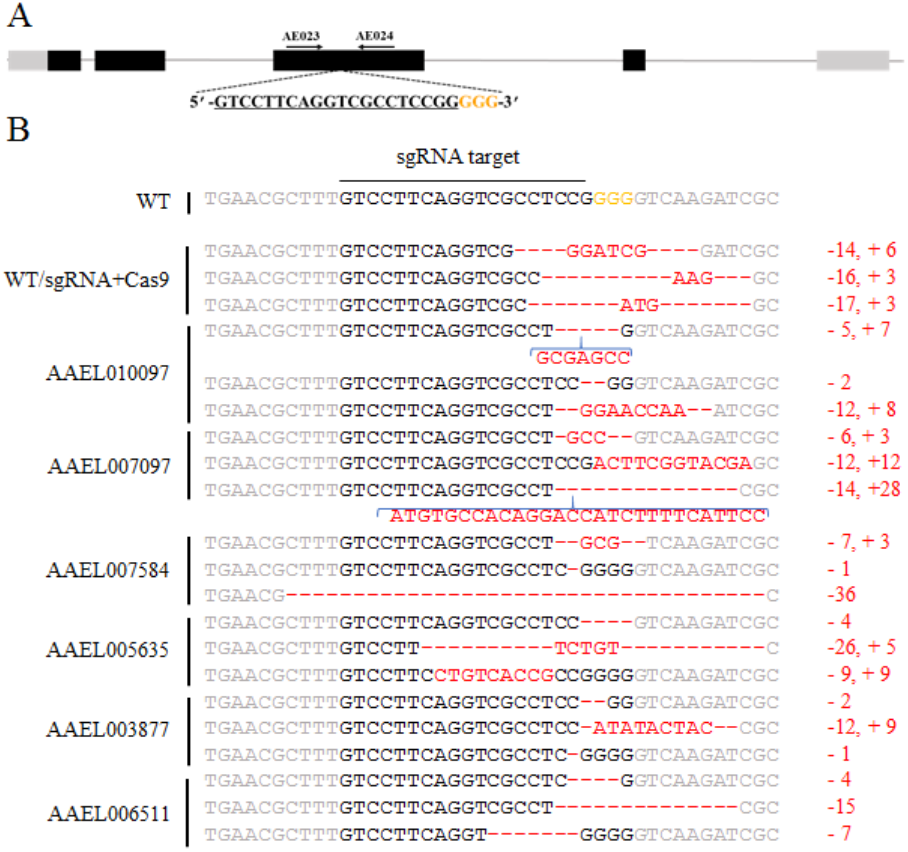
Confirmed mutagenesis of the *white* locus. Schematic representation of the *white* locus with exons indicated as boxes, coding regions depicted in black, and the 5’and 3’UTR’s in gray. Locations and sequences of the sgRNA targets are indicated with the PAM highlighted in yellow (A). Genomic sequencing analysis of indels from individuals sequenced from the various Cas9 strain injections. Top line represents WT sequence; PAM sequences (NGG) are indicated in yellow, and *white* gene disruptions resulting from insertions/deletions are indicated in red (B).

**Supplementary Figure 5.**
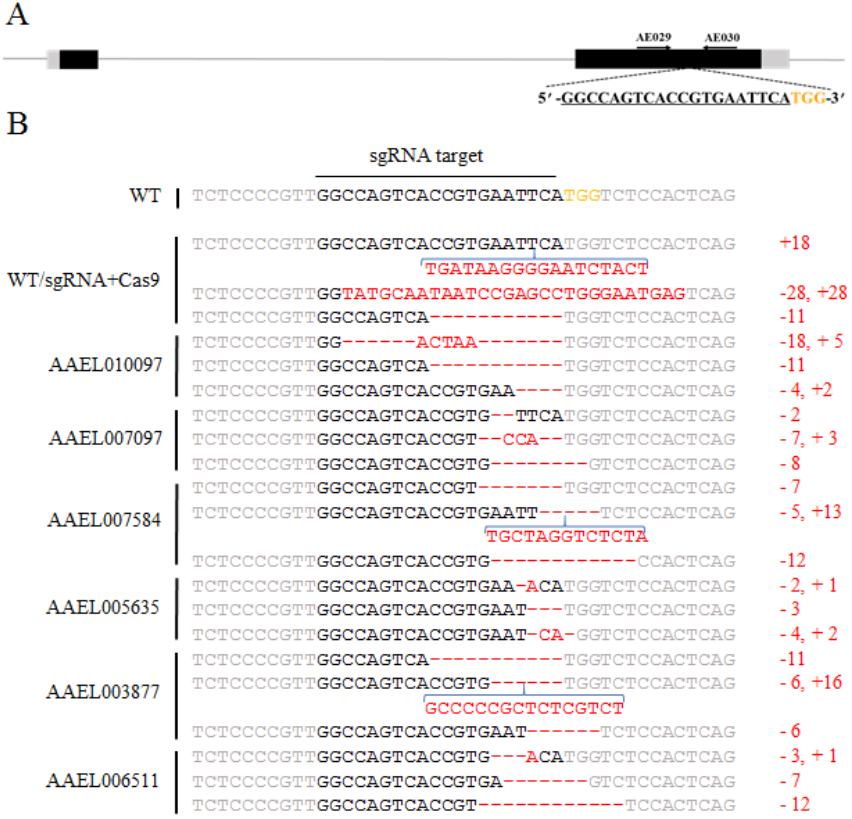
Confirmed mutagenesis of the *yellow* locus. Schematic representation of the *yellow* locus with exons indicated as boxes, coding regions depicted in black, and the 5’and 3’UTR’s in gray. Locations and sequences of the sgRNA targets are indicated with the PAM highlighted in yellow (A). Genomic sequencing analysis of indels from individuals sequenced from the various Cas9 strain injections. Top line represents WT sequence; PAM sequences (NGG) are indicated in yellow, and *yellow* gene disruptions resulting from insertions/deletions are indicated in red (B).

**Supplementary Figure 6.**
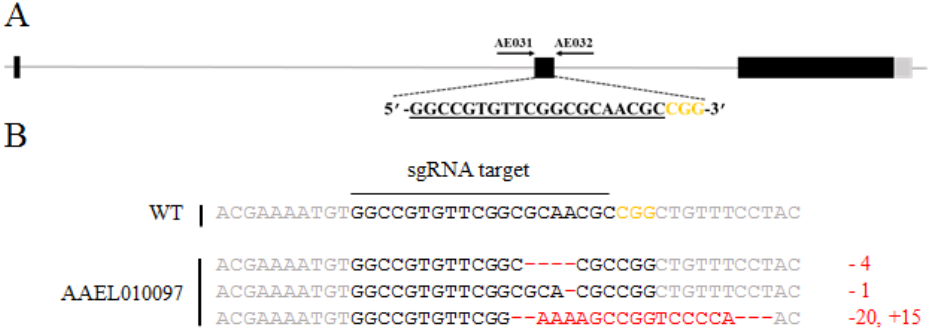
Confirmed mutagenesis of the *ebony* locus. Schematic representation of the *ebony* locus with exons indicated as boxes, coding regions depicted in black, and the 5’and 3’UTR’s in gray. Locations and sequences of the sgRNA targets are indicated with the PAM highlighted in yellow (A). Genomic sequencing confirms the generation of small indels (B). Top line represents WT sequence; PAM sequences (NGG) are indicated in yellow, and *ebony* gene disruptions resulting from insertions/deletions are indicated in red.

**Supplementary Figure 7.**
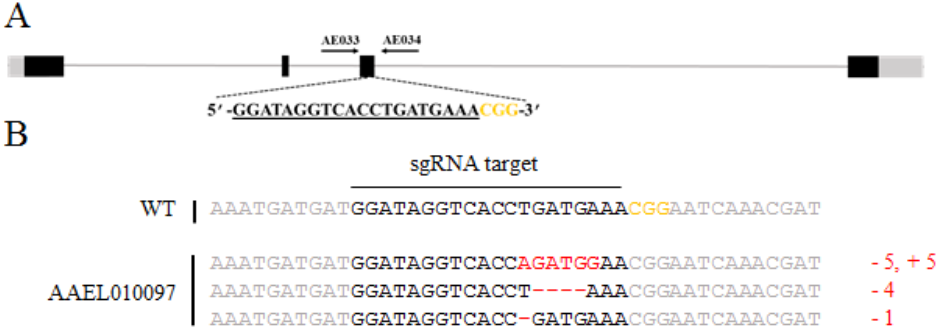
Confirmed mutagenesis of the *deformed* locus. Schematic epresentation of the *deformed* locus with exons indicated as boxes, coding regions depicted in black, and the 5’and 3’UTR’s in gray. Locations and sequences of the sgRNA targets are indicated with the PAM highlighted in yellow (A). Genomic sequencing confirms the generation of small indels (B). Top line represents WT sequence; PAM sequences (NGG) are indicated in yellow, and *deformed* gene disruptions resulting from insertions/deletions are indicated in red.

**Supplementary Figure 8.**
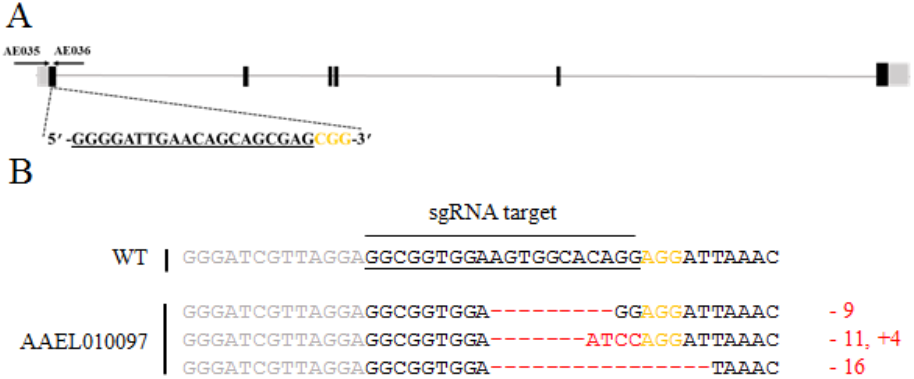
Confirmed mutagenesis of the *sine oculis* locus. Schematic representation of the *sine oculis* locus with exons indicated as boxes, coding regions depicted in black, and the 5’and 3’UTR’s in gray. Locations and sequences of the sgRNA targets are indicated with the PAM highlighted in yellow (A). Genomic sequencing confirms the generation of small indels (B). Top line represents WT sequence; PAM sequences (NGG) are indicated in yellow, and *sine oculis* gene disruptions resulting from insertions/deletions are indicated in red.

**Supplementary Figure 9.**
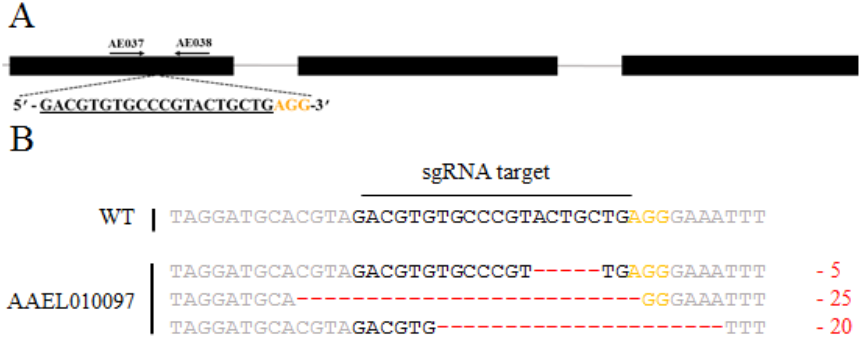
Confirmed mutagenesis of the *vestigial* locus. Schematic representation of the *vestigial* locus with exons indicated as boxes, coding regions depicted in black, and the 5’and 3’UTR’s in gray. Locations and sequences of the sgRNA targets are indicated with the PAM highlighted in yellow (A). Genomic sequencing confirms the generation of small indels (B). Top line represents WT sequence; PAM sequences (NGG) are indicated in yellow, and *vestigial* gene disruptions resulting from insertions/deletions are indicated in red.

**Supplementary Figure 10.**
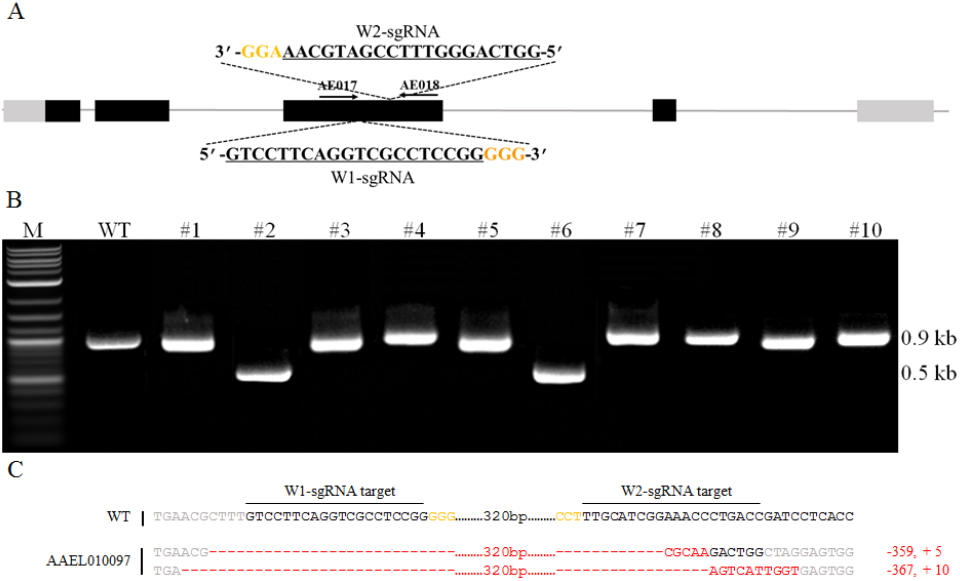
Generation of indels in the *white* locus by simultaneous injection of multiple sgRNAs. Schematic representation of the *white* locus with exons indicated as boxes, coding regions depicted in black, and the 5’and 3’UTR’s in gray. Locations and sequences of the sgRNA targets are indicated with the PAM highlighted in yellow. PCR and gel analysis of the *white* locus following simultaneous injection of two sgRNAs into the AAEL010097-Cas9 strain. In total 10 independent injected G0’s were tested along with wild type (WT) as a control.Generation of indels (0.4kb) for mosquitoes #2 and #6 can be straightforwardly distinguished on a DNA gel (B). Sequencing analysis of these indels confirms large deletions (C). Top line represents WT sequence; PAM sequences (NGG) are indicated in orange, and *white* gene disruptions resulting from insertions/deletions are indicated in red.

**Supplementary Figure 11.**
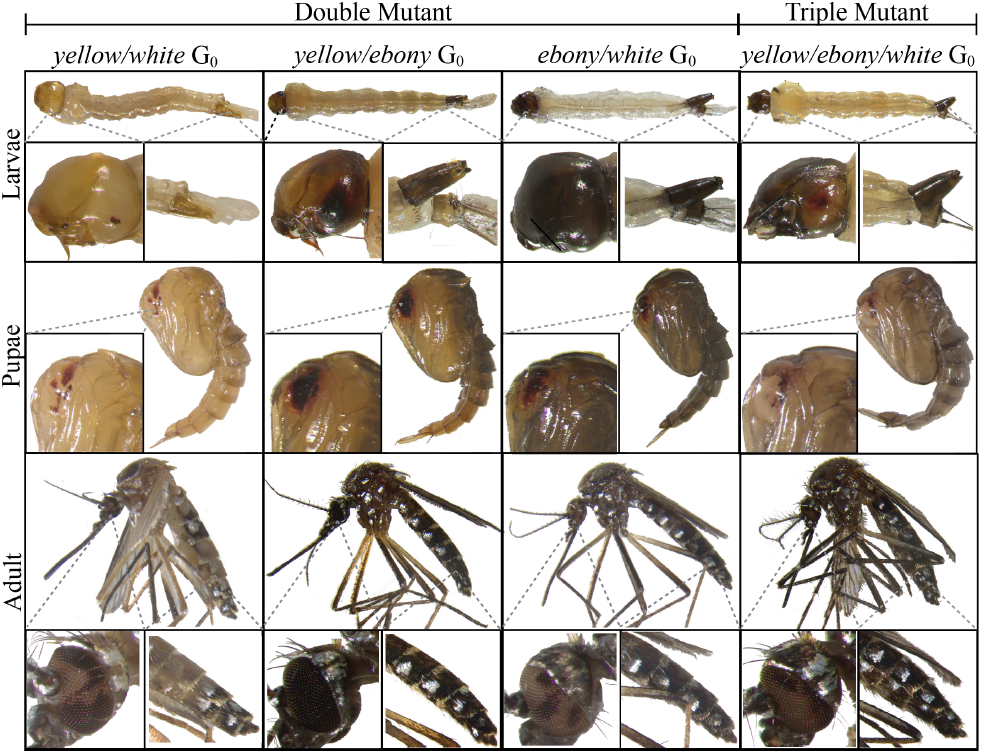
CRISPR/Cas9 induced double and triple G0 mosaic mutant phenotypes from single injections. Larval, pupa and adult G0 mosaic phenotypes of double mutants including yellow body and white eyes (*yellow/white)*, a mixture of yellow and dark body (*yellow/ebony)*, dark body and white eyes (*ebony/white)*, and one triple mutant which is a phenotypic mixture of yellow and dark body and white eyes (*yellow/ebony/white)*.

**Supplementary Figure 12.**
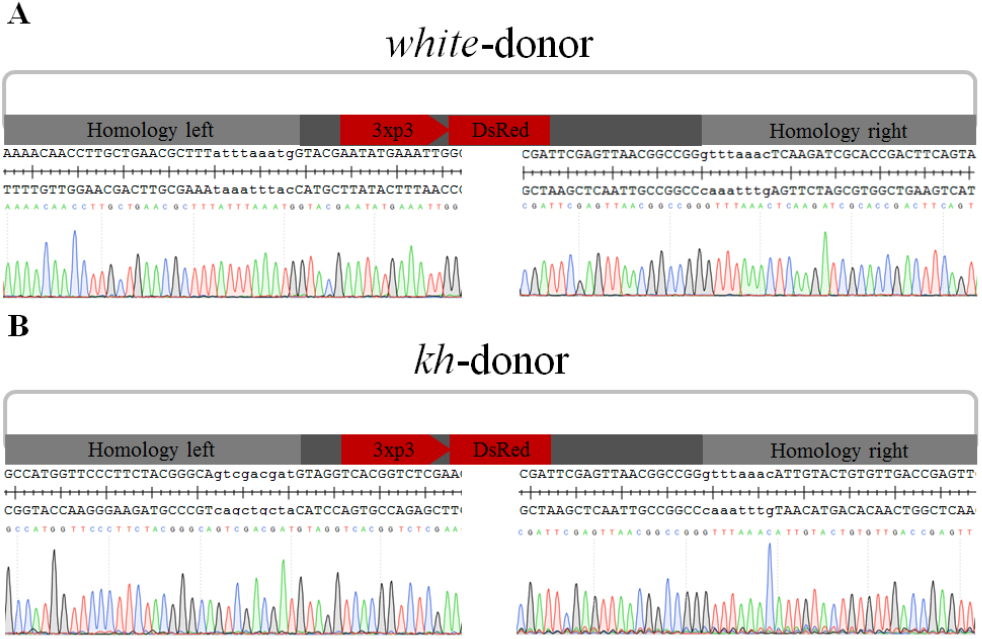
Confirmation of the precise donor insertions in the target loci via HDR. Genomic DNA sequencing was performed to confirm the donor construct insertions in *white* and *kh*. Sequencing of the 3’-end amplification fragments with primers of AE025 and AE026 (left), and 5’-end amplification fragments with primer AE027 and AE028 (right) from the *white* donor G1 founder (A). Sequencing of the 3’-end amplification fragments with primers of AE019 and AE020 (left), and 5’-end amplification fragments with primer AE021 and AE022 (right) from the G1 founder (B). Together, these results reveal precise integration of the *white*-donor and *kh*-donor construct into the target locus.

**Supplementary Table 1.**
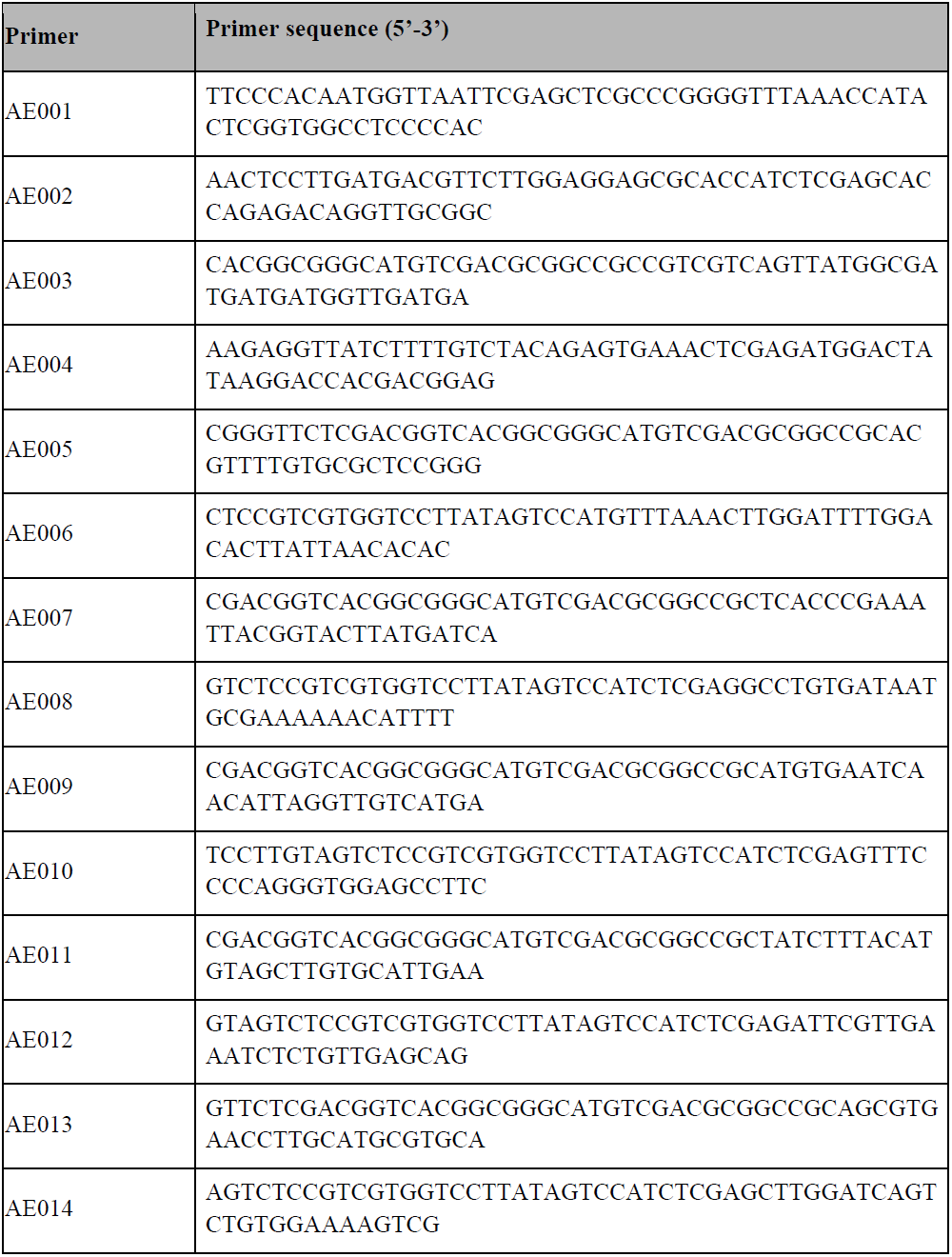
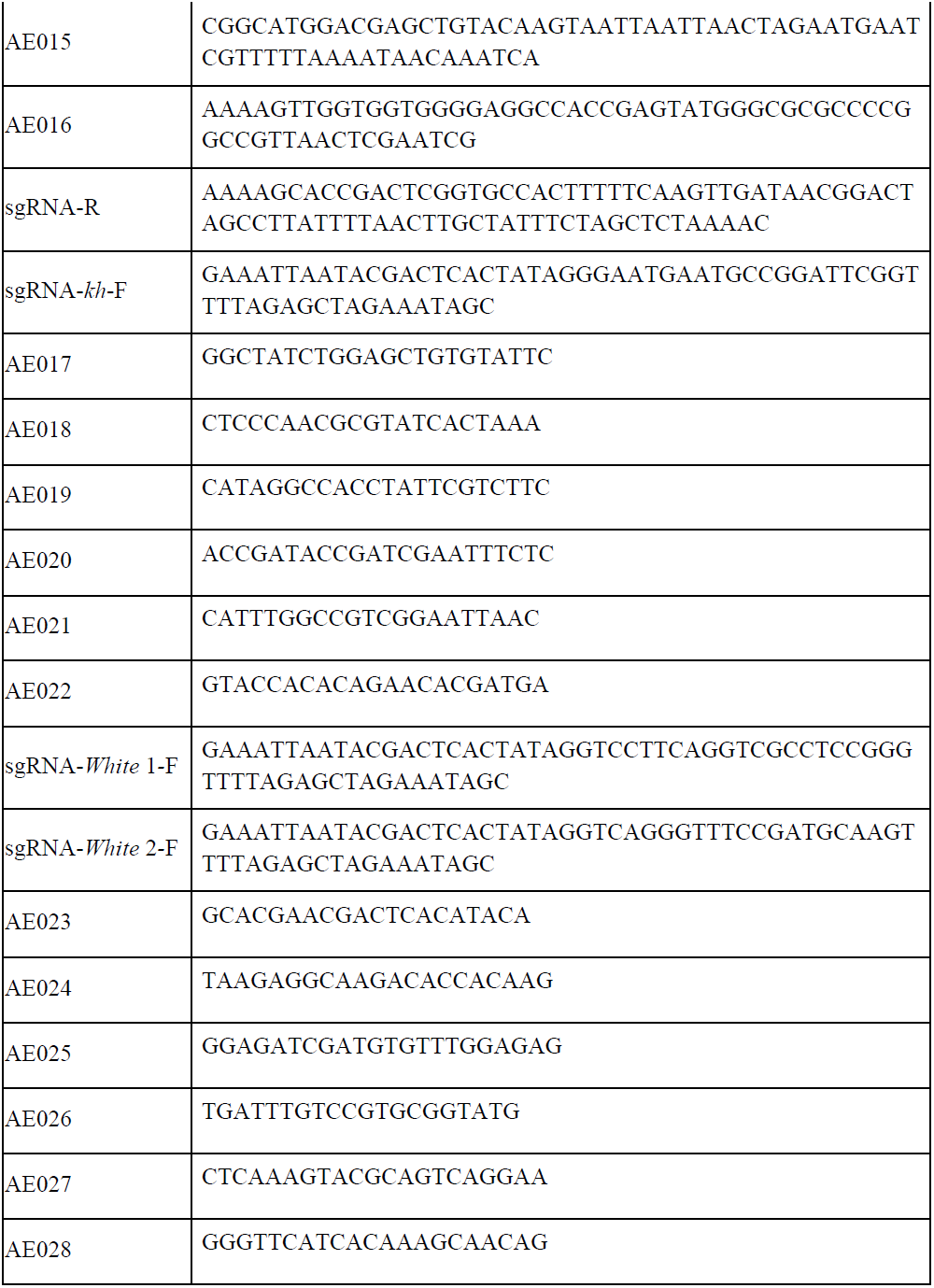
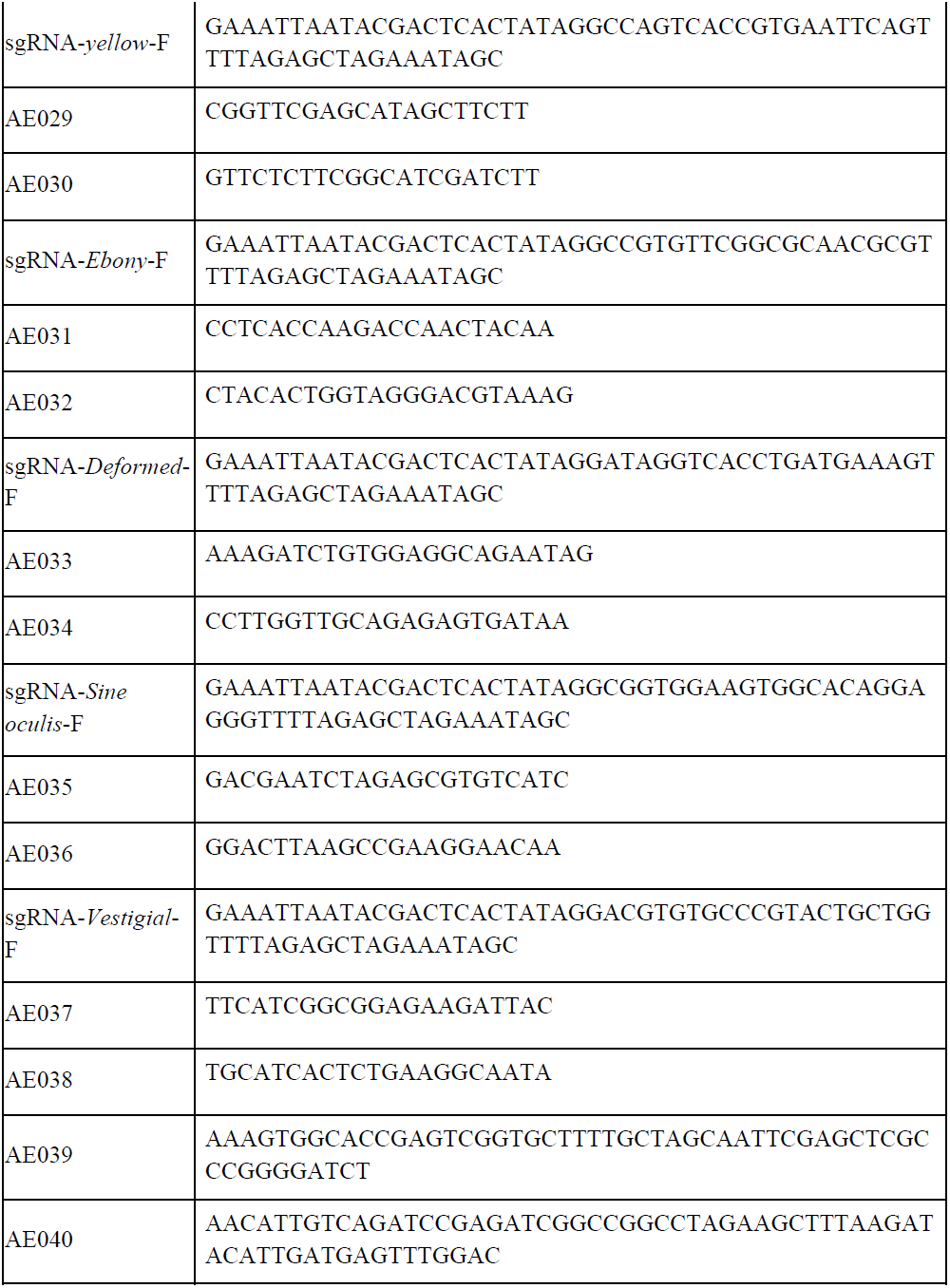
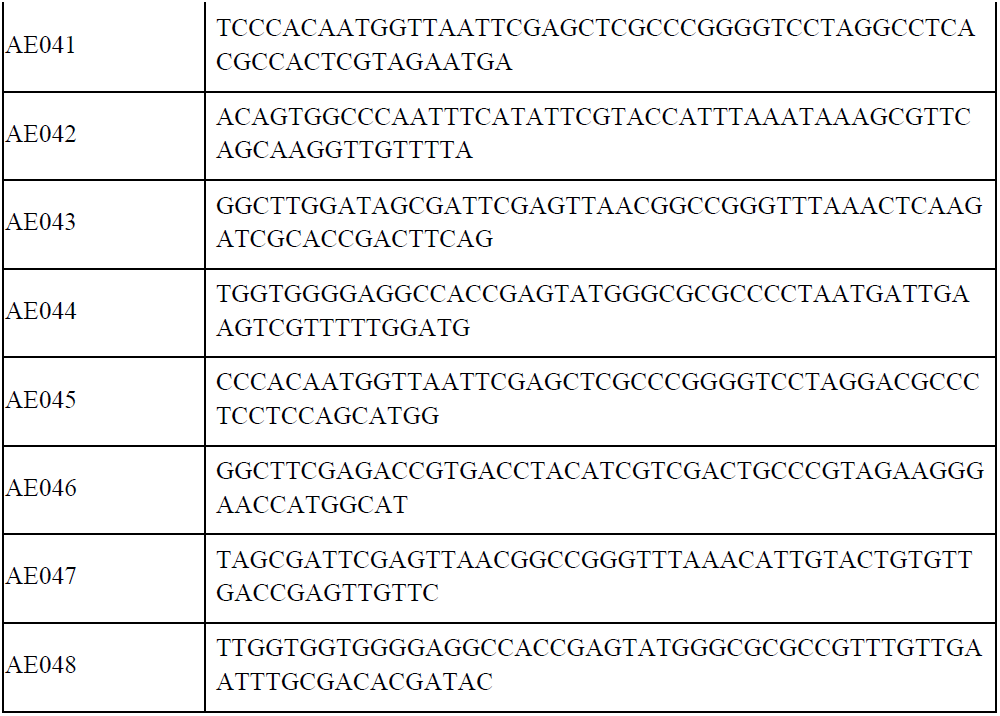
Supplementary table of primer sequences used in this study

